# Molecular characterization and culture optimization of marine copepod *Oithona dissimilis* (Landberg, 1940) from Nagore coastal waters, Southern India

**DOI:** 10.1101/2024.02.14.580272

**Authors:** P. Raju, P. Santhanam, B. Balaji Prasath, M. Divya, R. Prathiviraj, S. Gunabal, P. Perumal

## Abstract

This study focused on identifying the cyclopoid copepod *Oithona dissimilis* found in the Nagore coastal waters of Southern India. Based on both morphological and molecular approaches to identify this species. Morphological characters were confirmed by examining the arrangement of setae and spines on the exopod of swimming legs 1-4. Molecular studies were performed using the CO 1 gene, which proved an effective marker for species identification. After identifying the *Oithona dissimilis*, reared them under laboratory conditions to determine the effects of various environmental parameters on their survival, nauplii production, and population density. Tested different temperatures, light intensity, pH, salinity, and diets and found that the optimum conditions for rearing *Oithona dissimilis* under laboratory conditions were a salinity of 25 PSU, a temperature range of 24-28° C, pH of 8, light intensity of 500 lux, and a mixed algal diet at a concentration of 30,000 cells/ml. The present study confirms the importance of accurate taxonomic identification for the *Oithona* group at the species level. Additionally, our findings show that rearing cyclopoid copepods under laboratory conditions that mimic their natural range of environmental parameters is crucial for the thriving culture of these organisms, the aquaculture industry frequently utilizes this as a live feed.

**Graphical abstract:** 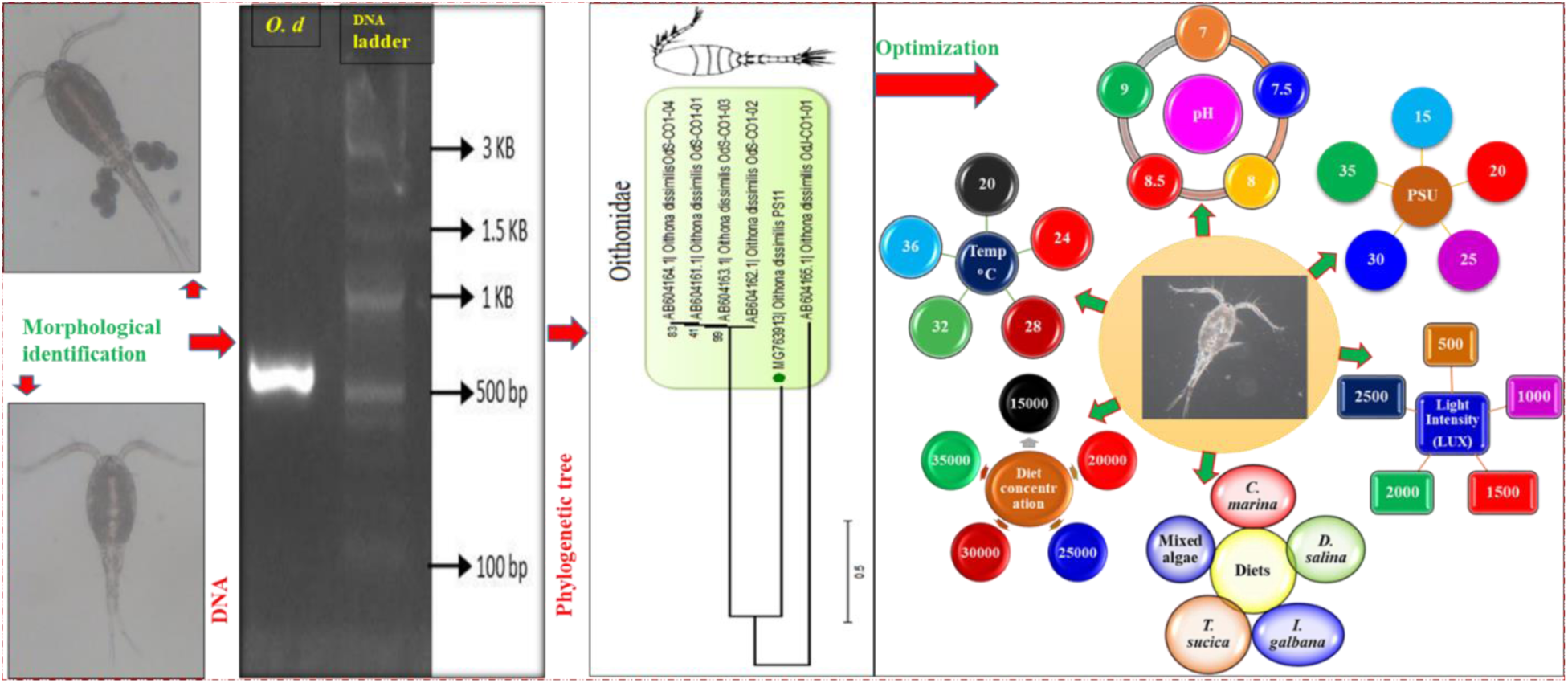

## INTRODUCTION

Many predatory fish in pelagic environments rely on tiny copepods as their primary food source, which are more plentiful than any other live-feed organisms (Spinelli et al., 2011; Shansudin et al., 1997; Ajiboye et al., 2011). The crustacean genus, *Oithona* is represented by small-size pelagic copepods that are distributed all over the world’s oceans and seas particularly in tropical and polar seas (Dvoretsky and Dvoretsky, 2015; Paffenhöfer, 1993; Saiz et al., 2003; Wang et al., 2015 Chew and Chong, 2011; Nielsen and Andersen, 2002; Chew et al., 2015). In the tropical region, these forms are dominantly distributed in neritic areas (Chew and Chong, 2011; Rezai et al., 2004). Being microscopic forms, they play an important role in the regeneration and exporting of nutrients (McKinnon and Ayukai, 1996, Zamora-Terol et al., 2014a). *Oithona*, which is a type of copepod, has a very important role in the marine food chain as it acts as a link between different species. The copepod feeds on various things such as phytoplankton and microbial components. However, the copepods are not safe from predators as they are being hunted by larger zooplankton and several pelagic ichthyoplankton (Spinelli et al., 2011; Castro et al., 2010; Van Noord et al., 2013). Despite their richness and important ecological role in the function of tropical marine diversity, only very little information is available for the *Oithona* group, especially on their biology and ecology.

The copepod, *Oithona dissimilis* Lindberg, 1940 is a copepod that is prevalent in estuaries located in the South East Asian continent and Islands of the tropical and subtropical West Pacific. This species is a prominent member of the zooplankton community and can be found widely distributed across these regions (Ferrari, 1977; Oka and Saisho, 1994; Lo et al., 2004; Saitoh et al., 2011). It’s been found that identifying the species of the genus *Oithona* can be quite challenging due to their small body size and subtle morphological differences. However, scientists have recently been using molecular identification techniques to more accurately identify and classify these copepods. This approach has proven to be quite effective and has helped us learn more about these fascinating creatures. There is a lack of information regarding cyclopoid copepods, particularly in relation to *Oithona* species. So far, studies have been concentrated on *O. similis, O. atlantica*, *O. nana* (Georgina et al., 2012) and *Dioithona rigida* (Radhika et al., al 2017). It’s important to note that there is currently no available data for the species *Oithona dissimilis*, which creates some uncertainty in the taxonomic classification of the Oithona group at the species level. One solution to this problem is to use a “total evidence” approach, which combines both morphological and molecular evidence to identify copepods. This method has been effective in the past (Mcmanus and Katz, 2009). It’s worth noting that copepods are an important food source for many fish and crustaceans, and using them as a live feed can boost larval survival and growth rate due to their high HUFA content and a broad range of body size. It’s interesting to note that while live feed such as Artemia and Rotifer are widely utilized in aquaculture for larval rearing practices, they may not provide all the essential nutrients required for optimal growth and development. As a result, there has been much research focused on mass culturing copepods to provide a more complete and nutritious food source for larvae. Various research laboratories around the world are currently working on developing copepod cultures that could potentially be used in the aquaculture industry. (Santhanam and Perumal 2011). However, due to inconsistency in production due to inefficient culture procedures, the copepods have not become popularized among aquafarmers. It is quite challenging to standardize the growth and reproduction of cyclopoid copepods in the field. However, to determine the ideal requirements for production, growth, and reproductive parameters, conducting laboratory experiments on the culture of copepods using natural environmental parameters is the best approach. Such an experiment is crucial for successfully cultivating copepods that can serve as live feed in the aquaculture industry (Hernandez Molejon and Alvarez-Lajonchere, 2003; James and Al-Khars, 1986). Temperature, light intensity; salinity, pH, diets and diet concentration have significant effects on the physiology and developmental stages of copepods. It is noteworthy that the cultivation of copepods is contingent upon key factors such as temperature, salinity, and diet. These variables hold considerable sway over the population density of the cultured copepods. It is therefore logical that the present investigation is focused on the collection, morphological and molecular characterization of the cyclopoid copepod, *Oithona dissimilis*, and the optimization of culture parameters for its production. Such efforts have the potential to yield a more comprehensive and nourishing food source for larvae in the aquaculture domain.

## MATERIALS AND METHODS

### Sample collection and identification

Samples of zooplankton were gathered from the Nagore coastal waters (as seen in Fig. 1) (Lat. 10.83° 03ˈ N; Long. 79.86° 47ˈ E) using a plankton net made of bolting silk cloth (No. 10, mesh size 158-µm) for approximately 20 minutes in the early morning. The samples were then immediately taken to the laboratory and vigorously aerated with a battery aerator. To reduce contamination of another zoo and meroplankton, the zooplankton sample was thoroughly rinsed. To isolate the size fractions that mainly contained adult and later-stage copepods, the zooplankter sample was screened. Rotifers, nauplii of copepod, and barnacles were eliminated by rinsing the samples through a zooplankton washer fitted with a 190 µm mesh size. To eliminate fish and prawn larvae, a first-course screening through a 500-μm mesh was performed. Specimens of the target cyclopoid copepod *Oithona* were then isolated and separated, and their morphological characteristics were observed using standard keys (Davis, 1955; Kasturirangan, 1963; Perumal et al., 1998 and Santhanam and Perumal, 2008). The isolated copepods were then observed under a stereo-phase contrast microscope and photographed with a digital still camera. After morphological identification, the separated copepods were preserved in 5% formalin for further morphological taxonomic study and 95% ethanol preservation for molecular analysis.

**Fig. 1.**
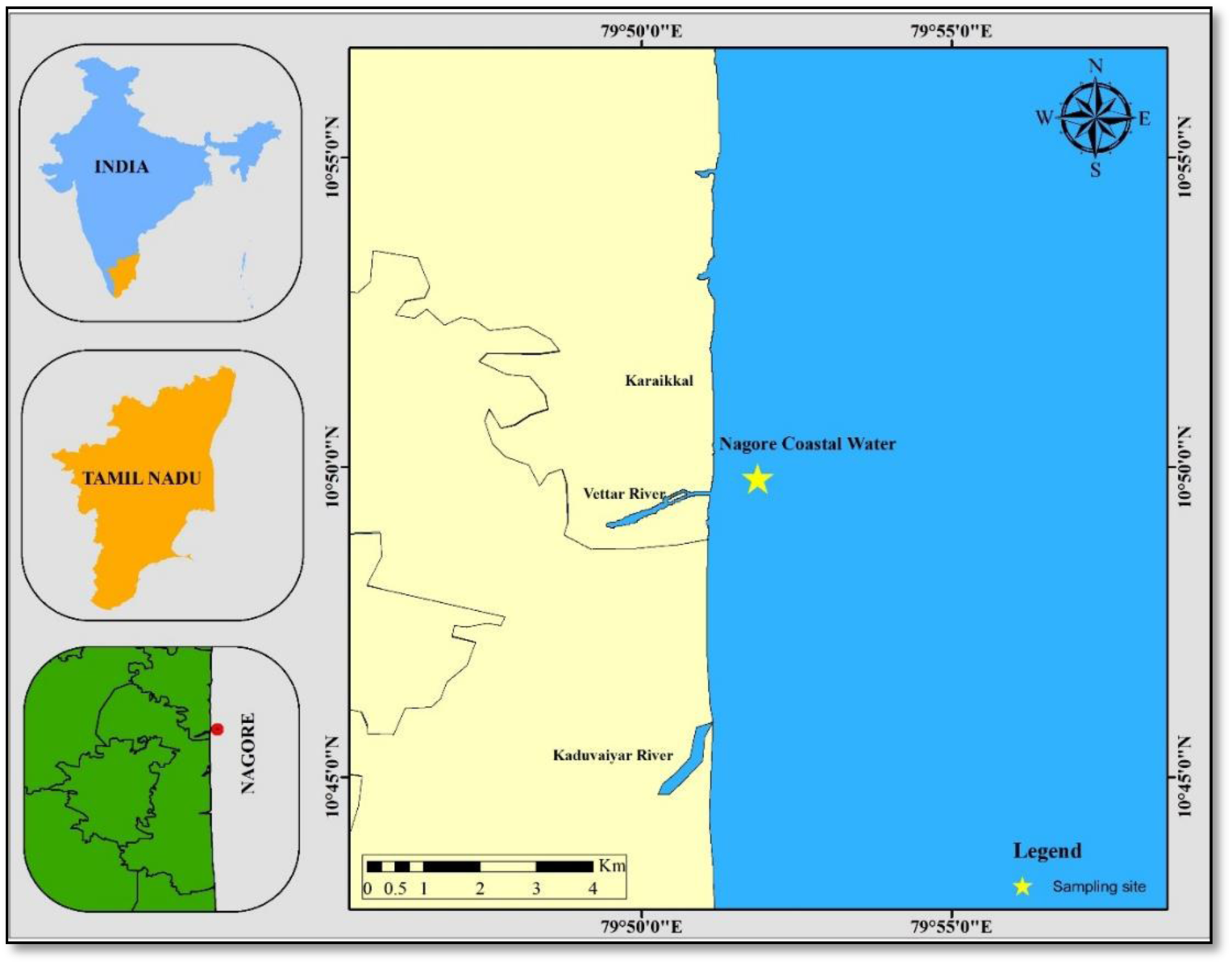
Map of the Nagore coastal waters showing the collection site.

### Genomic DNA isolation, PCR analysis and DNA sequencing

The Qiagen DNeasy tissue kit protocol was used to extract genomic DNA from *Oithona dissimilis*, a copepod that was identified. To perform the Polymerase Chain Reaction (PCR), 1 ml (50ng) of template DNA was mixed with 2ml (10pmol) of each Cytochrome c oxidase subunit I (CO1) primers that target LCO1490: 5′GGTCAACAAATCATAAAGATATTGG3′ and HC02198: 5′TAAACTTCAGGGTGACCAAAAAATCA-3′ (Folmer et al., 1994). The 20 ml mix consisted of 10 ml of PCR Master mix (Amplicon) and 5 ml of double-distilled water. The CO1 amplification process involved an initial denaturation at 94° C for 5 min, 30 cycles at 94° C (60 s), 52° C (60 s), and 72° C (60 s) followed by a final elongation at 72° C for 3 min before being transferred to 48°C until further analysis. An agarose gel electrophoresis (1.5%) was carried out to validate the PCR products. ACME Progen Biotech Pvt. Ltd. (Salem, India) was responsible for DNA sequencing the amplified product, which was edited in the Gene tool and Bio-edit software before being submitted to GenBank.

### Bioinformatics analysis

The Gene tool and Bio-edit software packages were used initially to edit the sequences. After editing, the sequences were submitted to the NCBI database. For phylogenetic analysis, DNA homology searches were carried out using the BLASTN 2.2.24 programs at NCBI, and similarity sequences were retrieved. To determine the levels of differentiation between genera and species, a multiple alignment of all similarity sequences was done by Clustal W 2.1. The phylogeny analysis was performed using the neighbour joining (NJ) search with Kimura 2-parameter as a model, which was carried out using MEGA version 4.0.2. The tree was bootstrapped using 1000 sub-replicates. Similarly, MEGA Ver. 4.0.2 was used to calculate the pair-wise nucleotide distances among the obtained partial 18S rRNA sequence and out-groups, by using the Kimura 2-parameter (Tamura et al., 2007).

### Microalgal culture

The cultivation of marine microalgae, namely *Isochrysis galbana* (ISO), *Chlorella marina* (CHL), *Picochlorum maculatum* (PICO), *Nannochloropsis oculata* (NAN) and *Amphora subtropical* (AMS), was carried out with great success. The process was conducted at the microalgae culture facility located at Bharathidasan University in Tiruchirappalli, India. The microalgae were grown at a temperature range of 23°-25°, with a salinity level of 30 PSU, and a light intensity of 45-60 mmol photons/m^2^/sec, for a light/dark cycle of 12 hours each. All of the microalgal strains were cultured using Walne’s (1974) Conway’s medium, and the seawater utilized in the culture underwent filtration with a 1μm filter bag, followed by sterilization via an autoclave. The containers utilized for the algal culture were meticulously sterilized before use. The harvested microalgae in the exponential phase were subsequently utilized as feed for copepods.

### Maintenance of copepod stock culture

In order to preserve the original culture, 50 male and female specimens of *O. dissimilis* were isolated and placed in a 1-liter beaker containing filtered seawater. The copepods were given a daily diet of mixed microalgae, consisting of equal amounts of ISO, CHL, PICO, NAN, and AMS, at a concentration of 30,000 cells/ml. The culture medium’s salinity and temperature were adjusted to 26PSU and 28°-30°, respectively. Daily removal of fecal pellets and debris, and replacement with fresh filtered seawater, ensured optimal conditions. The water quality parameters were consistently monitored to maintain pH, salinity, and temperature. *O. dissimilis* has a generation time of 10-12 days under optimal conditions, with 6 nauplii and 6 copepodite stages, including the adult. The adult gravid female copepods were used to restart mass culture, and the axenic copepod culture was maintained under controlled conditions at the Marine Planktonology & Aquaculture Laboratory.

### Experimental setup

#### Estimation of Survival Rate (SR)

The present study involved the execution of experiments to investigate the survival rates of copepods in varying water quality and dietary conditions. The study was carried out over 15 days and involved the use of ten healthy gravid female (*O. dissimilis*) individuals. The culture was sustained in a 100-milliliter beaker, which contained sterile seawater that was filtered through a 1-micrometer filter bag. The individuals were enumerated daily, and any deceased individuals were removed from the beaker. The experiments were conducted in triplicate and extended over a total period of 15 days. The daily removal of debris and faecal materials was necessary to maintain the culture.

#### Determination of Nauplii Production Rate (NPR)

The evaluation of the nauplii production capacity of *O*. *dissimilis* has been completed through the systematic assessment of its response to various environmental factors including temperature, light intensity, pH, salinity, different diets, and diet concentration. A mature female with viable egg sacs was placed in a test tube filled with 25ml filtered seawater. The release of nauplii was monitored every one or two hours. Once the nauplii were released, the adult female was removed from the test tube. The nauplii were subsequently counted under a microscope. This experiment was conducted in triplicate to ensure accuracy, and the collected data will undergo statistical analysis to determine the mean ± SE values.

#### Assessment of Population density (PD)

The population density of *O. dissimilis* was examined under various conditions, including water quality, diet, temperature, light intensity, pH, salinity, diet concentration, and different diets. To begin with, 10 adult copepods were isolated and inoculated into each 500-ml beaker filled with filtered sterilized seawater. This setup was maintained in triplicate. After 15 days, the animals were harvested through a 48µm mesh and fixed with 5% formalin. Finally, different stages of copepods (nauplii, copepodites, and adults) were counted under the microscope to estimate their population densities.

#### Statistical Analysis

The data obtained on the survival rate (SR), nauplii production rate (NPR), and population density (PDR) of *O. dissimilis*, regarding temperature, light intensity, pH, salinity, different diets, and diet concentration, have been analyzed using one-way ANOVA. In case of finding significant differences (P<0.05), Tukey’s multiple comparisons test has been applied to determine the specific difference among treatments. The data are presented as Mean±SE.

## RESULTS

### Morphological description of *O. dissimilis*

Female: The metasome has four segments, with each segment having a pair of dorsal sensory hairs except for segment 2, which has two pairs. The exopod of P1-P4, excluding the terminal spine, has 1-1-3, 1-1-3, 1-1-3, 1-1-2 external spines, respectively, while the endopod of P1-P4 has 0-0-1, 0-0-1, 0-0-1, 0-0-1 external setae, respectively, and 1-1-5, 1-2-5, 1-2-5, 1-2-4 internal setae, respectively. The 5th thoracic segment doesn’t have any hairs on the posterior margin. The caudal rami are longer than the 5th thoracic segment, and the proportions of the urosomal segments are as follows: 12, 33, 14, 14, 13. P5 has a fine seta that is directed dorsally and one terminal seta. There is no ciliation on either of these P5 setae, and the terminal seta reaches almost the end of abdominal segments 1-2. Male: The A1 is twice geniculated, with the proximal geniculation surrounded by a sheath and the distal geniculation having a notch in the segment. The caudal rami are shorter than in females, and Si is very short and can only be seen. The proportions of the autosomal segment’s caudal rami are 19: 19: 16: 13: 10: 11: 13. The prosome is laterally located and has a very complex group of integument organs in an area comprising the posterior ventral part of cephalosome and posterior extension or flap of cephalosome overlapping the following segment.

### Molecular characterization of CO1 gene of *O. dissimilis*

#### Blast

The dataset was prepared for our target species *O. dissimilis* Contig-PS11 based upon a similarity search. We selected the species based on the identity (>79%) and above 98% of query coverage (Table 1).

**Table 1:** Dataset preparation of cytochrome c oxidase I of *O. dissimilis* PS-11 and its phylogenetic similarity using NCBI-BLAST.

| Accession | Organism | Haplotype | Query cover (%) | E-value | Identity (%) |
| --- | --- | --- | --- | --- | --- |
| AB604163.1 | *Oithona dissimilis* | OdS-CO1-03 | 98 | 3.00E-144 | 81 |
| AB604164.1 | *Oithona dissimilis* | OdS-CO1-04 | 98 | 9.00E-144 | 81 |
| AB604161.1 | *Oithona dissimilis* | OdS-CO1-01 | 98 | 4.00E-142 | 81 |
| AB604162.1 | *Oithona dissimilis* | OdS-CO1-02 | 98 | 5.00E-141 | 81 |
| AB604165.1 | *Oithona dissimilis* | OdJ-CO1-01 | 98 | 4.00E-117 | 79 |

### Phylogenetic tree

#### Estimation of inter-and intra-specific phylogeny

The Neighbour-Joining analysis method was used to infer the evolutionary history of the COI gene of *O. dissimilis* PS-11. The optimal tree with a sum of branch length equal to 4.48590762 is presented. The percentage of replicate trees wherein the associated taxa clustered together in the bootstrap test (1000 replicates) is shown alongside the branches. The tree is drawn to scale and the branch lengths are in the same units as the evolutionary distances used for inferring the phylogenetic tree. Here we have constructed the phylogenetic tree of both inter- and intra-specific organisms. The inter-specific phylogeny shows that the COI gene of *O. dissimilis* PS-11 has diverged from the strains of OdS-CO1-01, OdS-CO1-02, and OdS-CO1-03; OdS-CO1-04 and thus the present form has been identified as *O. dissimilis* and the OdJ-CO1-01 act as an ancestor for our target species (Fig 3). The overall mean distance was found in the range of 1.782 and it surely indicates that O. *dissimilis* PS-11 is involved in the positive evolution of the Darwinian test for inter-specific phylogeny level. Whereas the intra-specific phylogeny reveals that, the tree was classified into two major clades and four sister clades. The first clade consists of four families viz., *Paracalanidae, Clausocalanidae, Centropagidae* and *Pontellidae* that are grouped. Whereas the second clade consists of the Oithonidae family as shown in Fig 5. The overall mean distance was found in the range of 0.150 and it surely indicates that *O. dissimilis* PS-11 is involved in the neutral evolution of the Darwinian test and no changes have been found to occur during the evolutionary process of inter-specific phylogeny.

**Fig. 2.**
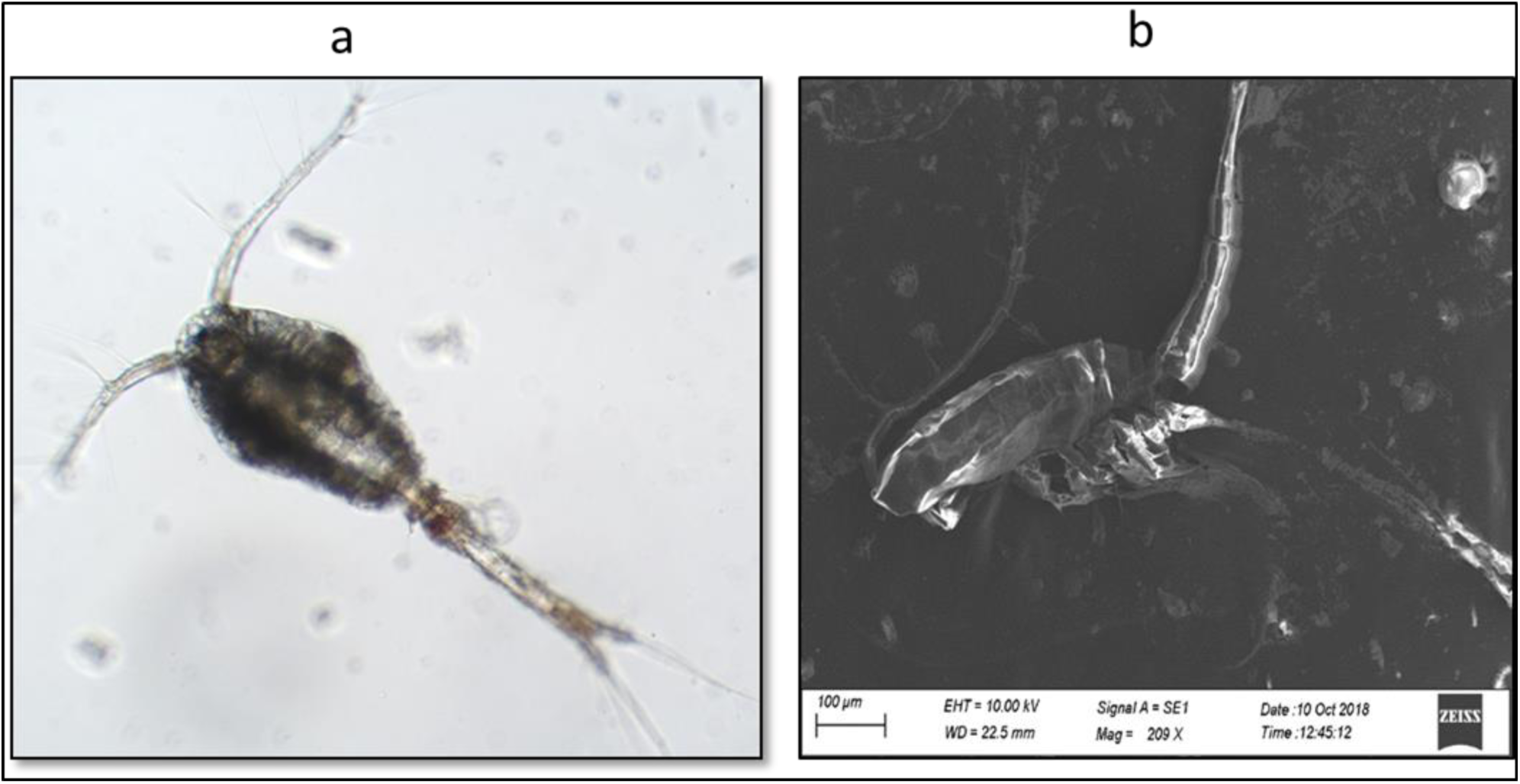
Stereo phase-contrast microscopic (a) and scanning electron microscopic (b) images of *O. dissimilis*.

**Fig. 3.**
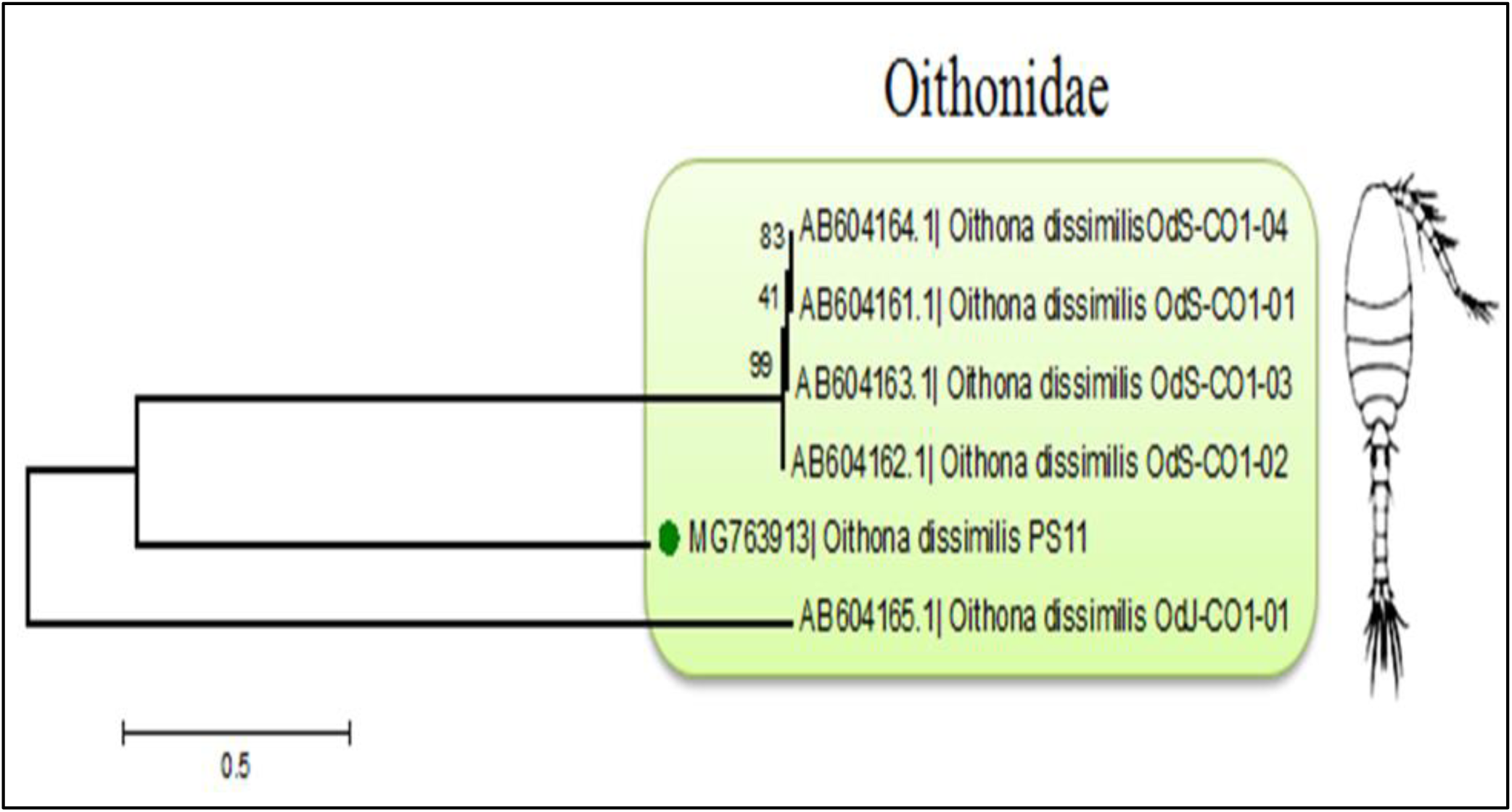
Construction of intera-specific phylogenetic tree of COI gene of *O. dissimilis* PS-11 from its closely related sequences obtained from MEGA 7.0. The green colour bullet differentiates our target sequences.

**Fig. 4.**
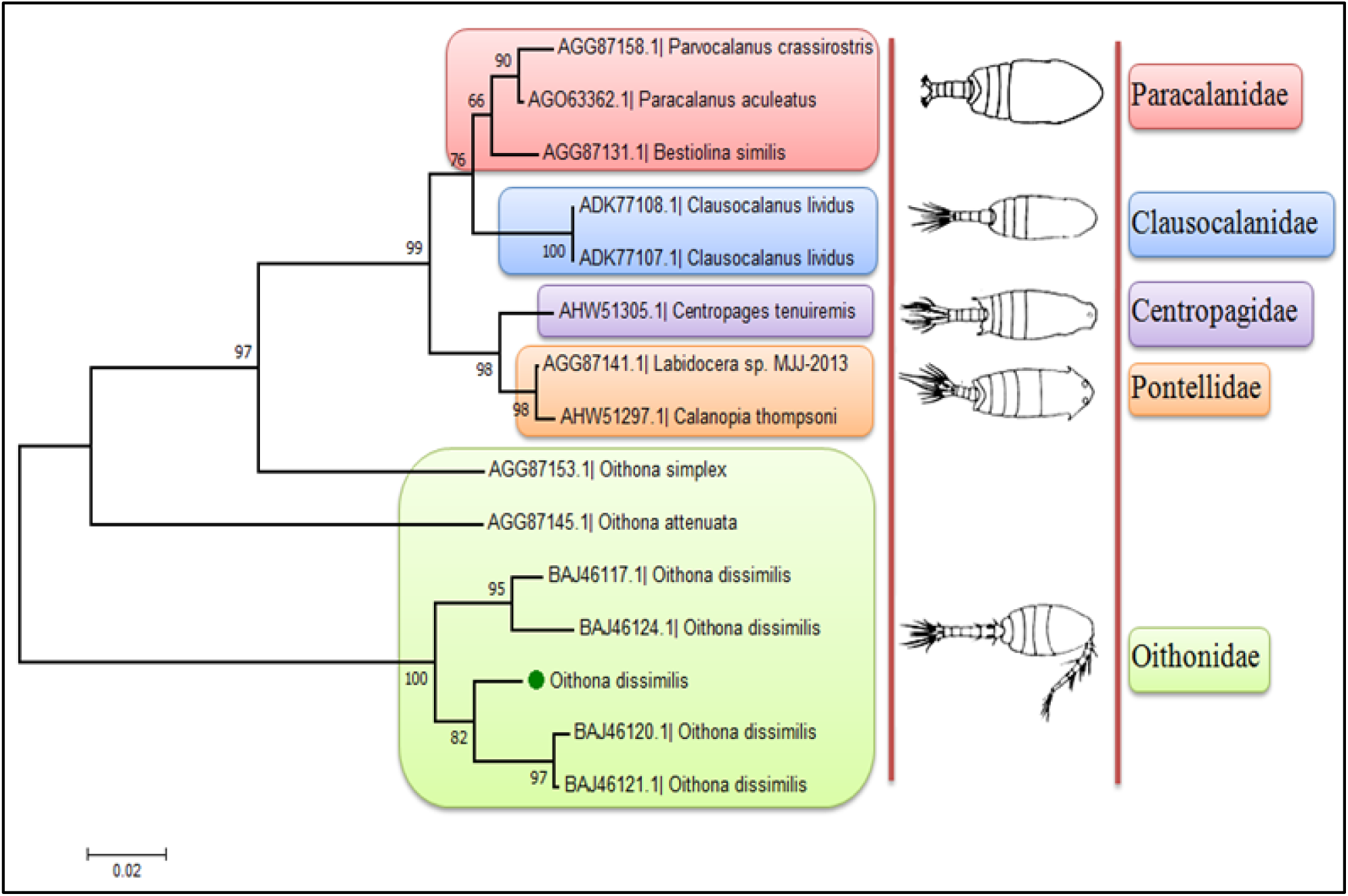
Construction of inter-specific protein-based phylogenetic tree of COI gene of *O. dissimilis* PS-11 from its closely related sequences obtained from MEGA 7.0. The green colour bullet differentiates our target sequence.

**Fig.5.**
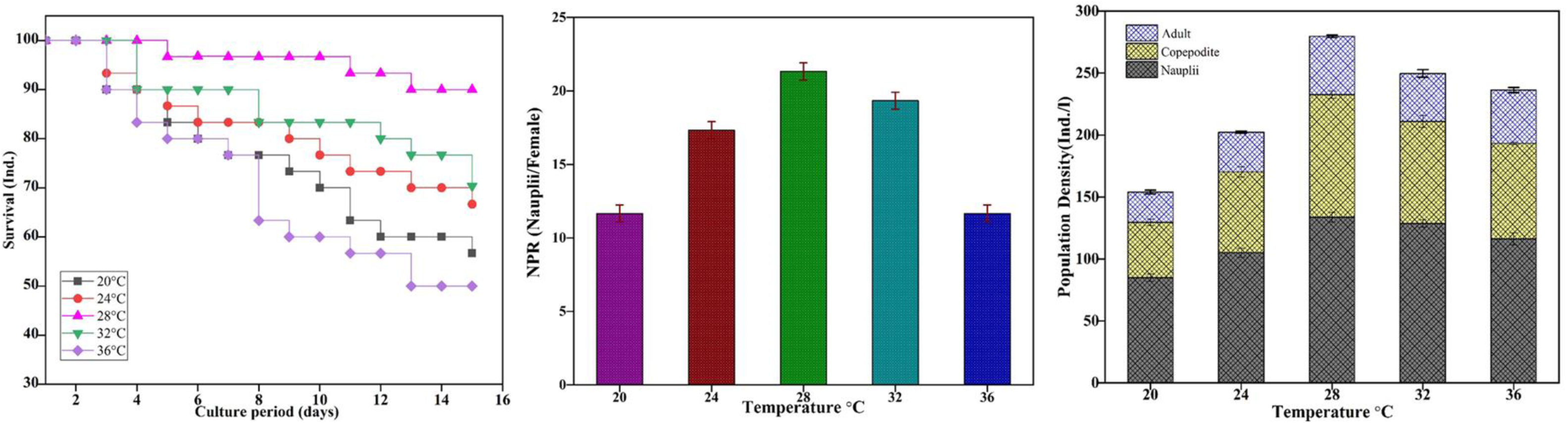
Effect of temperature on Survival rate (SR), Nauplii production rate (NPR) and Population density (PD) of *O. dissimilis*.

### Estimation of pair-wise genetic diversity

The maximum Composite Likelihood method was used to compute evolutionary distances in terms of the number of base substitutions per site. A total of 6 nucleotide sequences were analyzed, and any positions with gaps or missing data were eliminated. The genetic diversity of inter-specific phylogeny shows that *O. dissimilis* OdJ-CO1-01 is highly diverged when compared to the other strains of *O. dissimilis* and its occurrence range is between 1.031-1.137 (Table 2). Whereas, the intra-specific pair-wise genetic diversity shows that neutral evolution will take place and its diversity range was 0.0-0.274 (Table 3). This statistical pair-wise genetic diversity data shows that our study provided a strong conclusion.

**Table-2:**
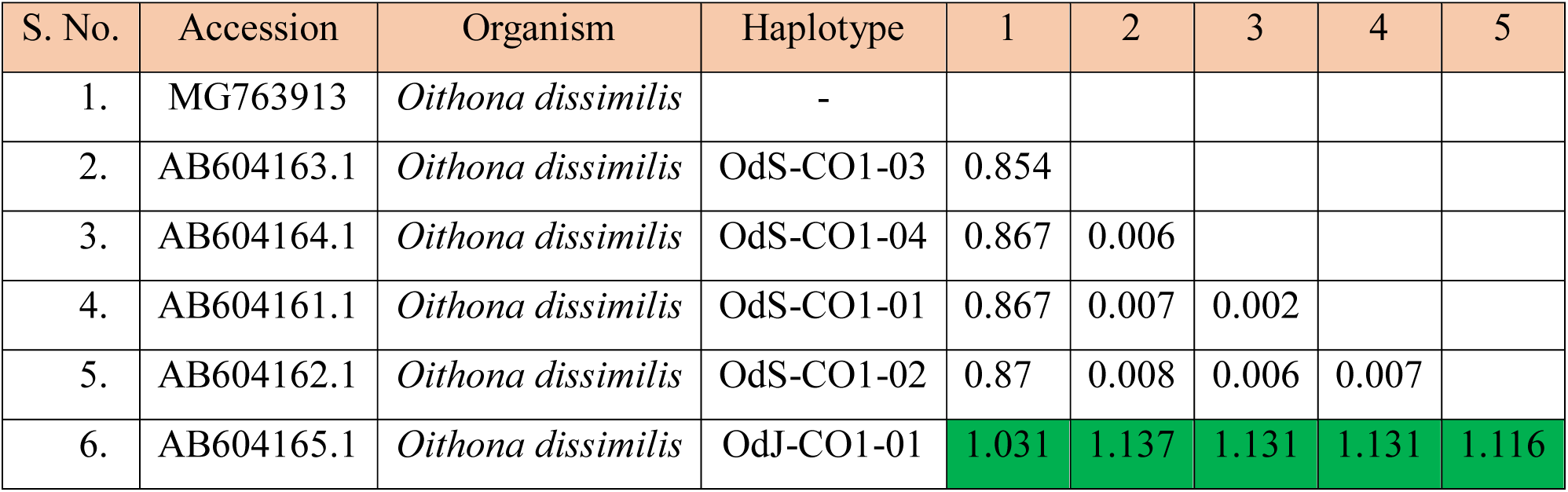
Estimation of inter-specific pair-wise genetic distance of COI gene of *O. dissimilis* PS-11 from its phylogenetic neighbours obtained from MEGA 7.0.

**Table-3:** Estimation of inter-specific pair-wise genetic distance of COI gene of O. dissimilis PS-11 from its phylogenetic neighbours’ obtained from MEGA 7.0.

| S. No. | Organism | 1 | 2 | 3 | 4 | 5 | 6 | 7 | 8 | 9 | 10 | 11 | 12 | 13 | 14 | 15 | 16 | 17 |
| --- | --- | --- | --- | --- | --- | --- | --- | --- | --- | --- | --- | --- | --- | --- | --- | --- | --- | --- |
| 1 | *Oithona dissimilis* |  |  |  |  |  |  |  |  |  |  |  |  |  |  |  |  |  |
| 2 | *Oithona dissimilis* | 0.032 |  |  |  |  |  |  |  |  |  |  |  |  |  |  |  |  |
| 3 | *Oithona dissimilis* | 0.037 | 0.005 |  |  |  |  |  |  |  |  |  |  |  |  |  |  |  |
| 4 | *Oithona dissimilis* | 0.046 | 0.051 | 0.056 |  |  |  |  |  |  |  |  |  |  |  |  |  |  |
| 5 | *Oithona dissimilis* | 0.065 | 0.065 | 0.07 | 0.023 |  |  |  |  |  |  |  |  |  |  |  |  |  |
| 6 | *Parvocalanus crassirostris* | 0.251 | 0.268 | 0.262 | 0.262 | 0.274 |  |  |  |  |  |  |  |  |  |  |  |  |
| 7 | *Paracalanus acueatus* | 0.251 | 0.268 | 0.262 | 0.262 | 0.274 | 0.009 |  |  |  |  |  |  |  |  |  |  |  |
| 8 | *Labidocera sp.* | 0.268 | 0.28 | 0.274 | 0.268 | 0.28 | 0.061 | 0.051 |  |  |  |  |  |  |  |  |  |  |
| 9 | *Calanopia Thompson* | 0.274 | 0.286 | 0.28 | 0.274 | 0.274 | 0.065 | 0.056 | 0.005 |  |  |  |  |  |  |  |  |  |
| 10 | *Oithona simplex* | 0.274 | 0.28 | 0.28 | 0.268 | 0.251 | 0.125 | 0.125 | 0.125 | 0.125 |  |  |  |  |  |  |  |  |
| 11 | *Centropages tenuiremis* | 0.262 | 0.274 | 0.268 | 0.262 | 0.274 | 0.065 | 0.056 | 0.023 | 0.028 | 0.135 |  |  |  |  |  |  |  |
| 12 | *Clausocalanus lividus* | 0.251 | 0.268 | 0.262 | 0.268 | 0.28 | 0.046 | 0.037 | 0.061 | 0.065 | 0.146 | 0.061 |  |  |  |  |  |  |
| 13 | *Bestiolina similis* | 0.251 | 0.274 | 0.268 | 0.268 | 0.28 | 0.028 | 0.018 | 0.056 | 0.061 | 0.125 | 0.061 | 0.042 |  |  |  |  |  |
| 14 | *Labidocera sp.* | 0.268 | 0.28 | 0.274 | 0.268 | 0.28 | 0.061 | 0.051 | 0.0 | 0.005 | 0.125 | 0.023 | 0.061 | 0.056 |  |  |  |  |
| 15 | *Bestiolina similis* | 0.251 | 0.274 | 0.268 | 0.268 | 0.28 | 0.28 | 0.018 | 0.056 | 0.061 | 0.125 | 0.061 | 0.042 | 0.0 | 0.056 |  |  |  |
| 16 | *Clausocalanus lividus* | 0.251 | 0.268 | 0.262 | 0.268 | 0.28 | 0.046 | 0.037 | 0.061 | 0.065 | 0.146 | 0.061 | 0.0 | 0.042 | 0.061 | 0.042 |  |  |
| 17 | *Centropage abdominalis* | 0.268 | 0.268 | 0.262 | 0.268 | 0.28 | 0.065 | 0.056 | 0.018 | 0.023 | 0.125 | 0.042 | 0.046 | 0.065 | 0.018 | 0.065 | 0.046 |  |

### Effect of temperature on Survival rate (SR), Nauplii production rate (NPR) and Population density (PD) of *O. dissimilis*

Population density, nauplii production and survival rates were evaluated at various temperatures. It was noticed that there was a significant difference found in survival rate in the percentage of copepods. There was above 50 % of survival occurred in almost all the temperatures tested. The highest survival rate (90%) was found at 28° C on the final day which was followed by 32° C (70.33%), 24° C (66.6%) and 20° C (56%). whereas, the lowest survival rate (50%) was observed at 36° C. However, there was a gradual decrease in survival found from the beginning of the first day towards the final day at a temperature of 36° C (Fig. 5). In all the test trials performed, the temperature was found to affect the nauplii production rate. The highest NPR (21.33 nauplii/female) was recovered at 24° C which was significantly higher (P<0.001) than at 20° C as well as at 36° C except at 24° C and 32° C which showed a considerably significant difference (P<0.05). The lowest NPR was found at 20° C and 36° C with only 11.66 nauplii/female which was significantly lower (P<0.001) when compared to the other temperatures tested (Fig. 5).

In all the trials conducted, the total highest population density (279.7 D/L) was obtained at 28° C which was significantly higher (P<0.001) than the rest of the temperatures tested. The lowest population density (154 D/L) was obtained at 20° C which was significantly lower (P<0.001) as compared to the other treatments. Thus, during all the trials, the temperature significantly affected the population density at different life stages of the copepod (Fig.5).

### Effect of salinity on Survival rate (SR), Nauplii production rate (NPR) and Population density (PD) of *O. dissimilis*

During the salinity trial, no significant variation was found in the survival rate of copepods. Above 50 % of survival was found in all the salinity levels tested except at 35 PSU.

The highest survival rate (96.66%) was found at 25 PSU on the final day followed by 30 PSU (73.33%), 20 PSU (70%) and 15 PSU (63.3%). The lowest survival rate (40%) was obtained at 40 PSU salinity level. However, there was a gradual decline in survival from the beginning of the first day towards the final day at 40 PSU salinity level (Fig. 6). The salinity was found to affect the nauplii production rate. The highest NPR (22 nauplii/female) was observed at 25 PSU which was significantly higher (P<0.001) than at 35 PSU, 20 PSU (P<0.05), 15 PSU (P<0.01) and there was no significant difference (P>0.05) noticed with 30 PSU salinity level. The lowest NPR was found at 35 significantly lower PSU (P<0.001) than 25 PSU followed by 30 PSU (P<0. 01) except 15 PSU and 20 PSU which showed no significant difference (P>0.05) respectively (Fig.6).

**Fig.6.**
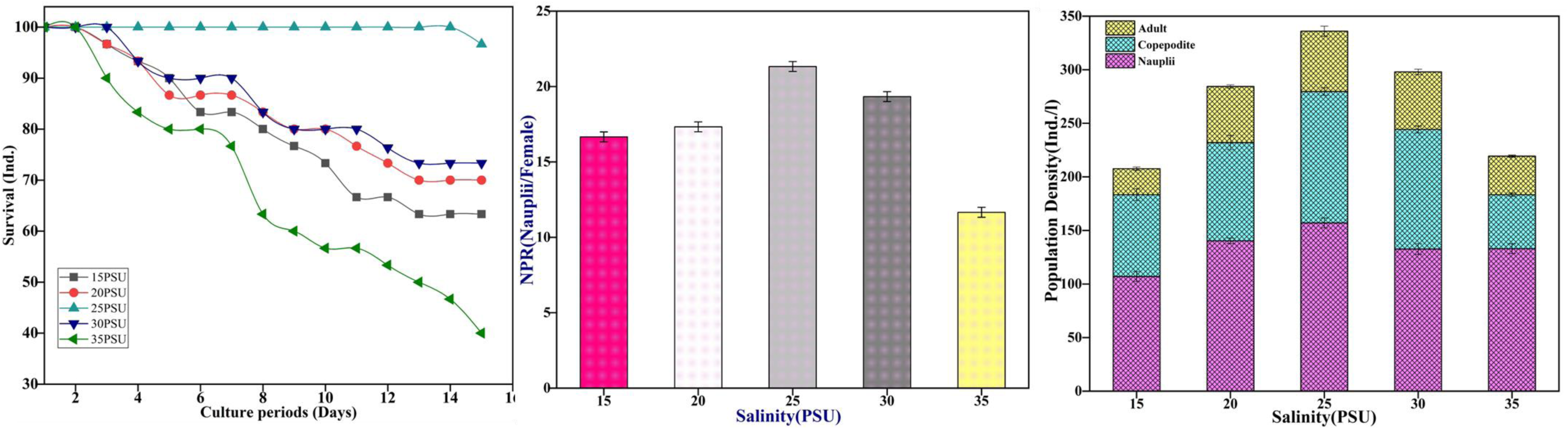
Effect of salinity on Survival rate (SR), Nauplii production rate (NPR) and Population density (PD) of *O. dissimilis*.

The highest total population density (336 ind. l) was obtained at 25 PSU and was significantly higher (P<0.001) than the rest of the salinities tested. The lowest population density (207.6 D/L) was obtained at 15 PSU which was significantly lower (P<0.001) when compared to the other treatments except at 35 PSU which did not show any significant difference (P>0.05). The same was the condition (P>0.05) in the population that occurred between 20 PSU and 30 PSU salinity levels. Thus, in all the trials the salinity significantly influenced the population density at different life stages. (Fig.6).

### Effect of pH on Survival rate (SR), Nauplii production rate (NPR) and Population density (PD) of *O. dissimilis*

In all pH trials conducted, there was above a 50% survival rate that occurred at all levels except at pH 7 and pH 9. In two cases of pH 7 and pH 9, there was a gradual decrease in the percentage of survival from the initial to final stage whereas the higher survival rate was noticed at pH 8 (86.6%), followed by pH 8.5 (83.3%). The lowest percentage of survival was found at pH 9 (40%) followed by pH 7 (43.3%) (Fig.7). The pH was found to affect the nauplii production rate in *O. dissimilis*. The highest NPR (19.66 nauplii/female) was found at pH 8 which was significantly greater (P<0.001) than at pH 7 (17.33 nauplii/female) and pH 9 (9.33 nauplii/female) followed by pH 7.5. There was no significant difference (P> 0.05) arising for pH 8 vs pH 8.5 and pH 7.5 vs pH 8.5 respectively. The lowest copepod nauplii production was recorded at pH 9 (9.33 nauplii/female) which was significantly lower (P<0.001) than with all other pH levels tested (Fig. 7). In the case of pH, the maximum population density (286.6 ind./l) was found at pH 8 which was greatly significance. The minimum density (94.6 ind./l) was observed at pH 7 which was significantly lower (P<0.001) when compared to other pH levels tested except at pH 9 which showed a considerable significant difference (P<0.05). Thus, in all the trials, pH significantly affected population density at different life stages. (Fig. 7).

**Fig.7.**
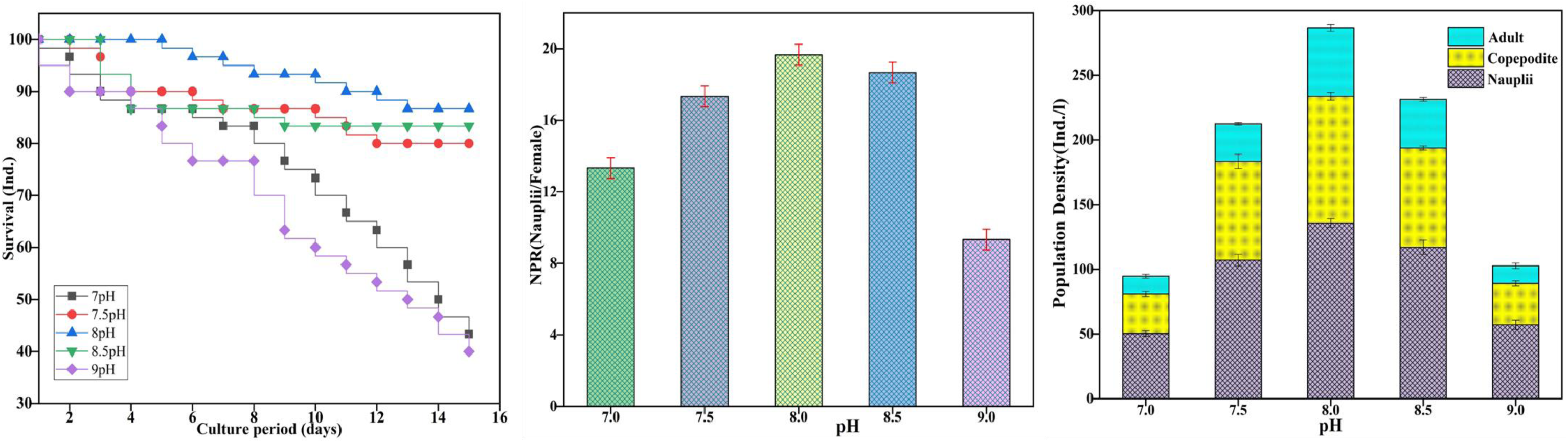
Effect of pH on Survival rate (SR), Nauplii production rate (NPR) and Population density (PD) of *O. dissimilis*.

### Effect of light intensity on Survival rate (SR), Nauplii production rate (NPR) and Population density (PD) of *O. dissimilis*

All the light intensity trials were tested and found that there was above 50% survival in almost all the intensities except at 2500 Lux where there was a lower survival rate (33.33%). In high intensity (2500 Lux) there was a gradual decrease in survival rate from the first day to the final day. The maximum survival (86.67%) was observed at low light intensity (500 Lux) followed by 1500 Lux (73.33%), 3000 Lux (63.33%) and 4500 Lux (53.33%) (Fig.8). The production rate of nauplii was affected by differences in light intensity. The maximum production (22.33 nauplii/female) was found at low light intensity (500 Lux) which was significantly greater (P<0.001) than 1000 Lux, 1500 Lux, 2500 Lux and 1000 Lux (P<0.01) respectively. The minimum production (8.66 nauplii/female) was found at higher light intensity (2500 Lux) which was significantly lower (P<0.001) than other intensities except for 2000 Lux which showed no significant difference (P>0.05). (Fig. 8). During culture at different light intensities, the total highest population density (276 ind. /l) was obtained at 500 Lux which was significantly higher (P<0.001) except 1000 Lux which did not show any significant difference (P>0.05). The lowest population density (180 ind./l) was obtained at 2500 Lux which was significantly lower (P<0.001) as compared to the other intensities tested except at 1000 Lux (P<0.01) and 2500 Lux (P<0.05). There was no significant difference noticed in population density under 1500 and 2000 Lux. Thus, in all the trials, light intensity significantly affected population density at different life stages. (Fig. 8).

**Fig.8.**
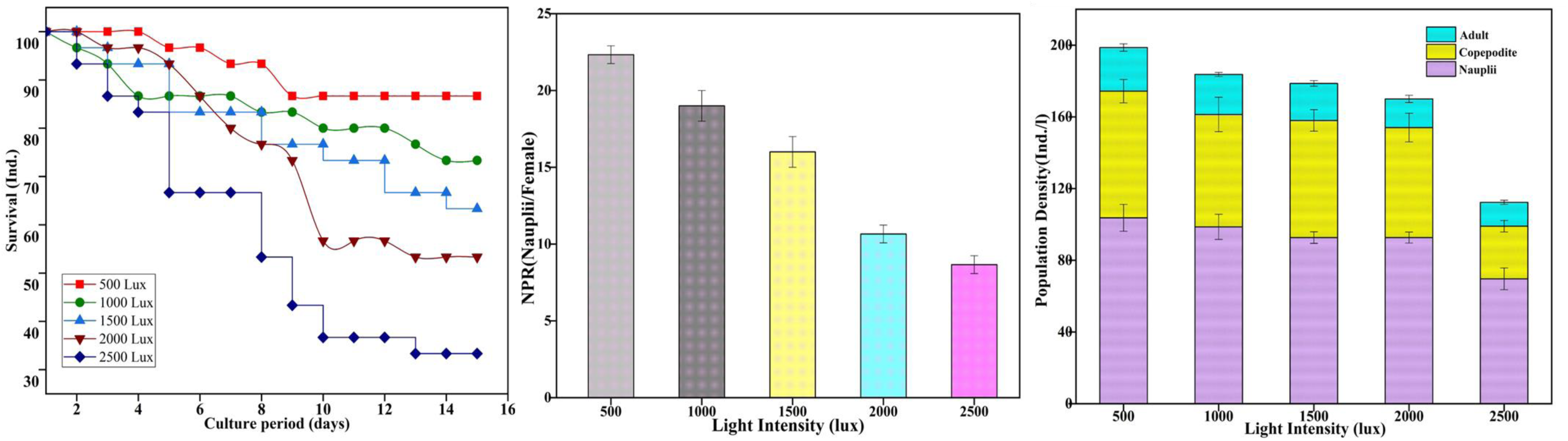
Effect of light intensity on Survival rate (SR), Nauplii production rate (NPR) and Population density (PD) of *O. dissimilis*.

### Effect of different feed on Survival rate (SR), Nauplii production rate (NPR) and Population density (PD) of *O. dissimilis*

In the case of different diet experiments, 50% of survival occurred in almost all algal feeds used however, the maximum survival rate (93.33%) was observed at *I. galbana* followed by *C. marina* (83.33%), mixed algae (76.67%) and *D. salina* (56.67%). The minimum survival rate (60%) was found at *T. suecica*. (9). The type of feed was found to influence the rate of production of nauplii in almost all the tests performed. The highest production rate (23.66 nauplii/female) was found at mixed algal feed which was significantly higher (P<0.001) than *T. suecica* followed by *C. marina* (P<0.05), *D. salina* (P<0.01) and for *I. galbana* (P>0.05) which showed no significant difference. The lowest production rate (16.33 nauplii/female) was noticed in the *T. suecica* diet which was significantly lower (P<0.001) than other feeds tested except *D. salina* which showed no considerable difference (P<0.05). (Fig. 9). At different algal feed experiments, the maximum peak (361.33 ind. /l) in population density was obtained in mixed algae which was significantly higher (P<0.001) than the rest of the feeds tested followed by *C. marina* (P<0.01) and I. galbana (P<0.05) respectively. The minimum population (267 ind./l) was obtained in copepod fed with *D. salina* which was significantly lower (P<0.001) compared to other feeds. There was no significant difference (P>0.05) found between *D. salina* and *T. suecica*. Thus, in all trials, different feed types significantly affect the population density of different life stages of the copepod (Fig. 9).

**Fig.9.**
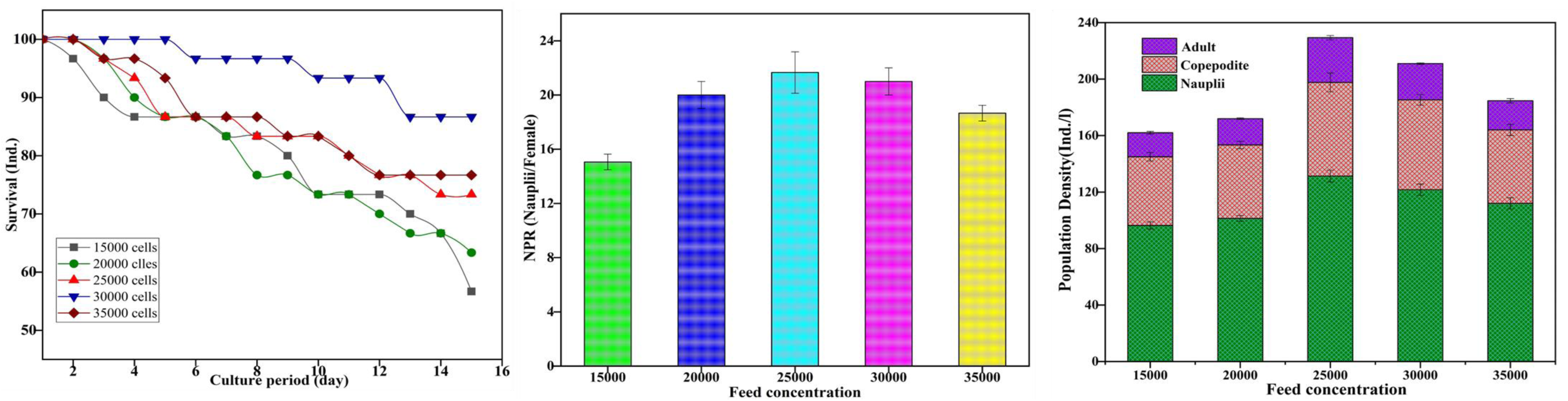
Effect of different microalgae concentration on Survival rate (SR), Nauplii production rate (NPR) and Population density (PD) of *O. dissimilis*.

### Effect of feed concentration on Survival rate (SR), Nauplii production rate (NPR) and Population density (PD) of *O. dissimilis*

During different concentrations of feed tested presently, there was 50% survival occurred in almost all the concentrations except 15000 cells/ml. There was a gradual decrease in survival percentage starting from the beginning and towards the end of the experiment at low concentrations (15000 cells/ml). The maximum survival of 86.67% was obtained at 30000 cells/ml followed by 76.67% at 35000 cells/ml, 73.33% at 25000 cells/ml and 63.33% at 20000 cells/ml. (10). Nauplii production rates were affected by different concentrations in feed. The maximum nauplii production rate (21.66 nauplii/female) was noticed at 25000 cells/ml which was significantly higher (P<0.001) than 15000 cells/ml followed by 35000 cells/ml (P<0.05) and there were no significant differences (P>0.05) in the concentration found with 30000 and 20000 cells/ml respectively. The lowest nauplii production (15.66 nauplii/female) was observed at low diet concentration (15000 cells/ml) which was significantly lower (P<0.001) than the other concentrations tested. (10).

At different feed concentrations, the total highest population density (229.3 D/L) was observed at 25000 cells/ml which was significantly higher (P<0.001) than the rest of the concentrations tested except at 30000 cells/ml which showed a considerable significant difference (P<0.05). The lowest population density (162 D/L) was found at 15000 cells/ml which was significantly lower (P<0.001) compared to other concentrations tested followed by 35000 cells/ml (P<0.01) and there was no significant difference (P<0.05) in population existed between 20000 and 15000 cells/ml. Thus, in all trials, concentration significantly affects population density at different life stages. (Fig.10).

**Fig.10.**
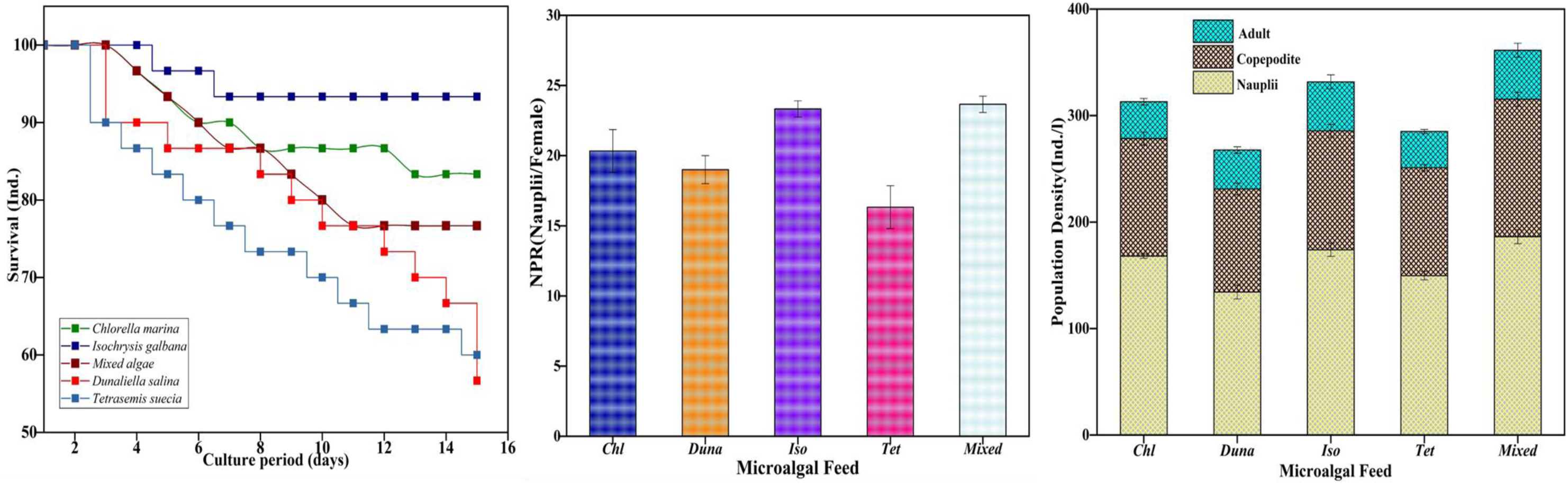
Effect of different microalgae feed on Survival rate (SR), Nauplii production rate (NPR) and Population density (PD) of *O. dissimilis*.

## DISCUSSION

*Oithona*, a cyclopoid copepod, is present in high numbers and holds significant ecological importance. This marine organism is extensively distributed throughout various marine environments. Identifying copepods of this specific genus on a regular basis is still difficult because of their petite size and inconspicuous morphological features that are used for diagnosis. (Radhika et. al., 2017). The copepod, identified in our study was very well characterized by the presence of the prominent features of *O. dissimilis* based on antenna (A1) which is shorter and both are geniculate in males and descriptive features confirmed that the males are usually smaller than females, urosome was 6 segmented in male and 5 segmented in female (Inshida 1985). To differentiate species within the genus, the prominent characters being conventionally followed are based on the arrangement of setae and spines on the exopod of swimming legs 1-4 (Radhika et al., 2017). Presently, the setae and spines of P1-P4 arranged in our specimen were consistent with the keys provided by Wellershaus (1969) and hence our copepod was identified as *O. dissimilis*. Bucklin et al. (2003) have confirmed that mt COI sequence variation has been proven to be a successful marker in molecular systematic and phylogenetic evolution in copepods. The utilization of the COI gene as the DNA barcodes has been proven to be an effective marker, especially for copepods (Hill et al., 2001; Bucklin et al., 2003). This gene has also been useful to distinguish the closely related genera for species identification (Paine et al., 2007). Accordingly, we have examined the mt CO 1 gene used for the identification and discrimination of *O. dissimilis* about the phylogenetic and evolution of copepods. Molecular phylogenetic analysis based upon mt CO1 revealed that our strain is distinct from the other related copepods. Presently, we have sequenced the mt CO 1 gene and compared its molecular features with the already publically available data on different species of different families available from NCBI. The mt CO1 gene of the species collected from Nagore coastal waters was subjected to BLAST and found that intra-species (*Oithona dissimilis*) was the most closely related species with 81% similarity (98% - Query coverage). Thus, our blast similarity was reliable with the finding of Soh et al., (2012) the individual who suggested that the CO 1 gene can be a suitable indicator for identifying species did so because it possesses sufficient variation to deal with both intra and inter-specific phylogenetic associations in invertebrates. (Soh et al., 2012). The phylogenetic relationships among *Oithona* sequences from NCBI with our selected samples using the mt COI gene were well resolved. CO 1 gene would be an appropriate biomarker for species discrimination as it has been widely employed to study population genetics and evolution (Shao and Barker, 2007). The mitochondrial genomes of animals contain a protein-coding gene that is the most conservative one found (Brown, 1985). It was found in our study that the overall mean distance for intra-specific phylogeny was found to occur in the range of 1.782 indicating that *O. dissimilis* PS-11 was involved for positive evolution of Darwinian test for intra-specific phylogeny level. For inter-specific phylogeny, the overall mean distance has occurred in the range of 0.150 indicating that, O. dissimilis PS-11 was involved in the Neutral evolution of the Darwinian test and no changes have occurred during the evolutionary process of inter-specific phylogeny. A higher level of genetic distance was found among intraspecies within our strain. The occurrence of higher levels of genetic distance in our strain with intra and inter-species levels might be due to the presence of cryptic or new species or sub-species and so on. Although the copepods have been shown to reveal higher levels of genetic divergence, sometimes the observed morphological conservatism might not follow the same level of genetic divergence. It is possible that the reason for this is the lack of separation between reproductive isolation and morphological divergence (Goetze 2003). Analysis of the CO 1 gene sequence has clearly demonstrated the occurrence of within-species variation in many crustaceans (Lefebvre et al., 2006). This variation is caused by the presence of cryptic or sibling species. The analysis has also identified similar levels of speciation in other eukaryotes (Waugh, 2007). Therefore, it is crucial to conduct detailed studies of the morphology, behaviour, and molecular characteristics of a population of closely related *Oithona* species in the future. In our present study, *O. dissimilis* was able to survive, and produce more nauplii and population density at a temperature range of 28°C - 32°C as reported by earlier researchers for other copepods (Rajthilak et al., 2014; Peter and Downing 1984; Kaviyarasan et al., 2019; Santhanam and Perumal, 2012).

*O. dissimilis* exhibited a high tolerance to salinity levels in various regions, according to our study. The salinity regions that the species was able to endure were wide-ranging of 25-30 PSU. This species belongs to the family Oithonidae and generally *Oithona* groups are commonly associated with coastal areas and are abundant in brackish water habitats. Our species was able to survive and produce nauplii and population density at the salinity range of 15-35 PSU. There was maximum mortality, low nauplii growth and low population density at low salinity (15 PSU) which might be due to the additional osmoregulation and respiration demands at these salinities (Kimoto et al., 1986; Santhanam 2012). The presently recorded maximum survival, nauplii production rate and total population density can be attributed towards the low light intensity of 500 Lux. The significant reduction in the production of offspring, as well as survival and growth rate of copepods, was observed with the increase in light intensity. Coping with the intense light conditions could have led to stress and energy consumption by copepods, which could be the possible mechanism for such a result. (Kaviyarasan et al., 2019; Farhadian, et al., 2014).

The development and fecundity of copepods are influenced by the quality of their food. Algal diets have a significant impact on the survival rate, nauplii production, and population of *O. dissimilis*. Our experiment confirmed that the copepods fed with mixed algae achieved the highest rate in terms of survival, nauplii production, and total population density. The reason for this could be that providing mono diets may lead to nutritional deficiencies in one or more vital nutrients. To minimize this risk, several researchers have suggested using mixed diets since the combined nutrient contents would fulfil the nutritional requirements of the target species (Brown et al., 1989; Santhanam and Perumal 2012; Smith et al., 1992). In response to feed concentration, the copepods had the highest survival rate, nauplii production and higher population density supplied with a higher concentration of algal cells but the ratio has declined in copepods supplied with a low concentration of algal cells (15000 cells/ml) might be due to food scarcity. Since food is one of the important factors in enhancing better growth and density of copepods in the culture systems, the copepod population increased in direct proportion to the increased food supply and poor results were obtained at low food concentrations (Schipp et al., 2009; Santhanam and Perumal 2012).

## CONCLUSION

Through a comprehensive study, we were able to successfully achieve the accurate identification of *O. dissimilis*. This involved a thorough examination of both morphological and molecular characteristics. Additionally, we were able to establish an optimization technique for commercial mass culture of this species under laboratory conditions. Our experimentation revealed that certain factors played a crucial role in the survival rates, population numbers, and nauplii hatching of *O. dissimilis*. Specifically, we found that a salinity level of 25 PSU, a temperature ranges of 24-28°C, a pH of 8, 500 lux of light, and a mixed diet with moderate to high concentration led to superior results for this particular species. Given these findings, we believe that *O. dissimilis* represents a viable live feed option for Aquaculture. Furthermore, the insights gleaned from our experiment could be utilized to develop an improved, commercial-scale copepod culturing system in the future. Overall, this study has important implications for the aquaculture industry and could contribute to the development of more sustainable and efficient practices.

## ACKNOWLEDGMENTS

The authors are extremely thankful to Bharathidasan University for providing the necessary facilities. The Ministry of Environment, Forest and Climate Change (MoEF & CC), Govt. of India, New Delhi, for providing financial support through a research Project MoEF & CC (F. No. 220180/06/2015-RE (Tax) 5th October 2016). One of the author (P. Raju) thank The Department of Biotechnology (DBT), Govt. of India, is greatly acknowledged for the copepod culture facility provided (BT/PR 5856/AAQ/3/598/2012).

## AUTHOR CONTRIBUTIONS

P. Raju: Data curation, methodology, Formal analysis, Writing-original draft; P. Santhanam: Conceptualization, Resources, Supervision, Funding, visualization, Writing-review & editing; B. Balaji Prasath: Data curation, Writing-review & editing; M. Divya: Formal analysis; R. Prathiviraj: Writing-review & editing; S. Gunabal: Formal analysis; and P. Perumal: Writing-review & editing.

## CONFLICT OF INTEREST STATEMENT

The authors declare no conflict of interest

## Notes

### Competing Interest Statement

The authors have declared no competing interest.

